# Higher baseline alpha power is associated with faster responses in visual search

**DOI:** 10.1101/2025.08.29.673162

**Authors:** Katharina Duecker, Kimron L. Shapiro, Simon Hanslmayr, Benjamin J. Griffiths, Andrew J. Quinn, Jeremy Wolfe, Yali Pan, Aleksandra Pastuszak, Ole Jensen

**Affiliations:** Department of Neuroscience, Brown University, RI, USA; Centre for Human Brain Health, School of Psychology, University of Birmingham, UK (current or former); School of Psychology and Neuroscience, Centre for Neurotechnology, University of Glasgow, UK; Brigham and Women’s Hospital, Boston, MA, USA; Harvard Medical School, Boston, MA, USA; School of Psychology, University of Nottingham, Nottingham, UK; Department of Experimental Psychology, University of Oxford, Oxford, UK; Oxford Centre for Human Brain Activity, Wellcome Centre for Integrative Neuroimaging, Department of Psychiatry, University of Oxford, Oxford, UK

## Abstract

Visual search models have long emphasised that task-relevant items must be prioritized for optimal performance. While it is known that search efficiency also benefits from active distractor inhibition, the underlying neuronal mechanisms are debated. Neuronal alpha oscillations (7-14 Hz) have been associated with functional inhibition of cortical excitability, as well as distractor suppression in spatial attention and visual working memory tasks. We therefore hypothesised that alpha oscillations similarly support the deselection of distractors in visual search. Using Magnetoencephalography (MEG), we here show that high alpha power before the onset of a complex search display is associated with faster search performance. Crucially, we used a General Linear Model (GLM) approach to control for confounds between alpha power and task duration, ruling out that this result was merely driven by practice effects paired with increased fatigue over time. In addition to spontaneous oscillatory activity, we quantified the cortical excitability to colours of the search stimuli based on Rapid Invisible Frequency Tagging (RIFT) responses. In contrast to our initial hypothesis, increased pre-search alpha power did not correlate with the RIFT response, providing no direct evidence for feature-specific inhibition of distracting stimuli by alpha. Our findings challenge the traditional view of alpha oscillations reducing visual processing, showing instead that increased occipital alpha power can enhance performance in a visual task. We propose that the increase in alpha power may reflect increased top-down control supporting visual search

## Introduction

Efficient search is the foundation of many daily activities, such as finding a friend in a crowd. This task is operationalised in visual search paradigms that require the rapid identification of a target stimulus among distractors. The attentional mechanisms underlying visual search are often conceptualized as a priority map, i.e. a representation of the visual field in which locations are weighted according to whether they are more or less likely to contain a target (Awh et al., 2012; Koch & Ullman, 1985; Navalpakkam & Itti, 2005; Serences & Yantis, 2006; Thompson & Bichot, 2005; Zelinsky & Bisley, 2015). Using Magnetoencephalography (MEG) and Rapid Invisible Frequency Tagging (RIFT), a subliminal visual stimulation technique, we have recently shown that a classic guided visual search task modulates neuronal excitability in early visual cortex in accordance with a model of the priority map (Duecker et al., 2025).

Here, we sought to test whether occipital alpha oscillations in visual cortex (8-12 Hz) benefit performance by facilitating the suppression of distractors, or whether they are detrimental to visual search.

A series of electrophysiological studies has demonstrated that alpha power is reliably modulated by attention (Gutteling et al., 2022; Hanslmayr et al., 2007; Kelly et al., 2006; Sauseng et al., 2005; Vissers et al., 2016; Worden et al., 2000). Indeed, alpha oscillations have been argued to support the suppression of distracting stimuli in visual tasks(Bonnefond & Jensen, 2024; Jensen & Mazaheri, 2010; Klimesch, 2012). Spatial attention paradigms, for instance, have repeatedly revealed an increase in alpha power over the cortical area processing distractors (Forschack et al., 2022; Foster et al., 2017; Kelly et al., 2006; Popov et al., 2019; Worden et al., 2000; Yuasa et al., 2023; Zhigalov et al., 2019), which has also been linked to the participants’ ability to ignore the stimulus (Gutteling et al., 2022; Händel et al., 2011; Händel & Jensen, 2014; Spaak et al., 2016; van Zoest et al., 2021; Zhao et al., 2023). Similarly, visual working memory tasks have revealed an increase in alpha power when several distractors have to be ignored (Bonnefond & Jensen, 2012; Jensen et al., 2002). This body of literature suggests that alpha oscillations may support visual tasks that require the deselection of irrelevant stimuli, such as visual search. For instance, alpha may aid visual search by selectively suppressing task-irrelevant features in a spatially unspecific way. Alternatively, alpha oscillations may serve as an inhibitory threshold applied to all stimuli. As target relevant features are boosted by feature-selective attention (Duecker et al., 2025; Maunsell & Treue, 2006), this mechanism could serve as a filter that only allows potentially task-relevant stimuli to pass, thus focusing the search on items that are likely to be a target (Ph.D. thesis Pastuszak, 2018). Currently, however, only pre-liminary findings suggest that high alpha power is associated with fast responses in complex visual search, where the locations of the target and distractors are unknown (Ph.D. thesis Pastuszak, 2018; Pastuszak et al., 2018).

Alpha oscillations have long been known to reflect functional inhibition of cortical activity as quantified by firing rates (Dougherty et al., 2017; Haegens et al., 2011), broadband high-frequency activity (Iemi et al., 2022), or the fMRI BOLD response (Scheeringa et al., 2011). After its initial discovery, the alpha rhythm was associated with inattention, boredom, and inactivation of the visual cortex (Adrian & Matthews, 1934; Berger, 1929; Foxe et al., 1998; Pfurtscheller et al., 1996). In line with that, signal detection tasks have repeatedly linked high alpha power to a higher number of misses (Dijk et al., 2008; Hanslmayr et al., 2007; Iemi et al., 2017). Based on this body of research, one might predict that high alpha power before the onset of the visual search display should be associated with worse search performance.

In this study, we investigated whether high alpha power is detrimental or beneficial to visual search. Moreover, we sought to test whether any correlation between alpha oscillations and search performance is associated with a modulation of visual responses to the search items. We therefore analysed MEG data from a visual search task, which required the identification of a single target among a high number of distractors (Figure 1, Duecker et al., 2025). Cortical responses to the different search stimuli were quantified using Rapid Invisible Frequency Tagging (RIFT, Duecker et al., 2025). Using a General Linear Model (GLM) approach accounting for confounding effects of time-on-task on alpha power (Benwell et al., 2019; Kopčanová et al., 2024) we show that higher alpha power is correlated with fast response times, without compromising hit rates. However, we were unable to link the increase in alpha power to reduced RIFT responses.

## Results

This study combines MEG and RIFT to study alpha oscillations and cortical excitability in complex visual search (see Duecker et al., 2025 for details). Participants (N=31) were instructed to indicate whether a single letter “T” was presented among 16 or 32 “L”s, without moving their gaze from the centre of the screen. In the *guided search* condition, a coloured T (either yellow or cyan) presented at the beginning of a block indicated the colour of the target for the duration of the block. A white T would signal the start of an *unguided search* block, during which the colour of the T was randomized over trials (Figure 1). Stimuli of the same colour were tagged at 60 or 67 Hz, by modulating their luminance sinusoidally. As we have reported previously, the RIFT response indicated a boosting of neuronal excitability to the known target colour and reduced responses to the distractor colour for guided compared to unguided search (Duecker et al., 2025). This modulation of the RIFT response as well as the participants’ task performance, are in line with priority-based mapping in early visual cortex.

**Figure 1.**
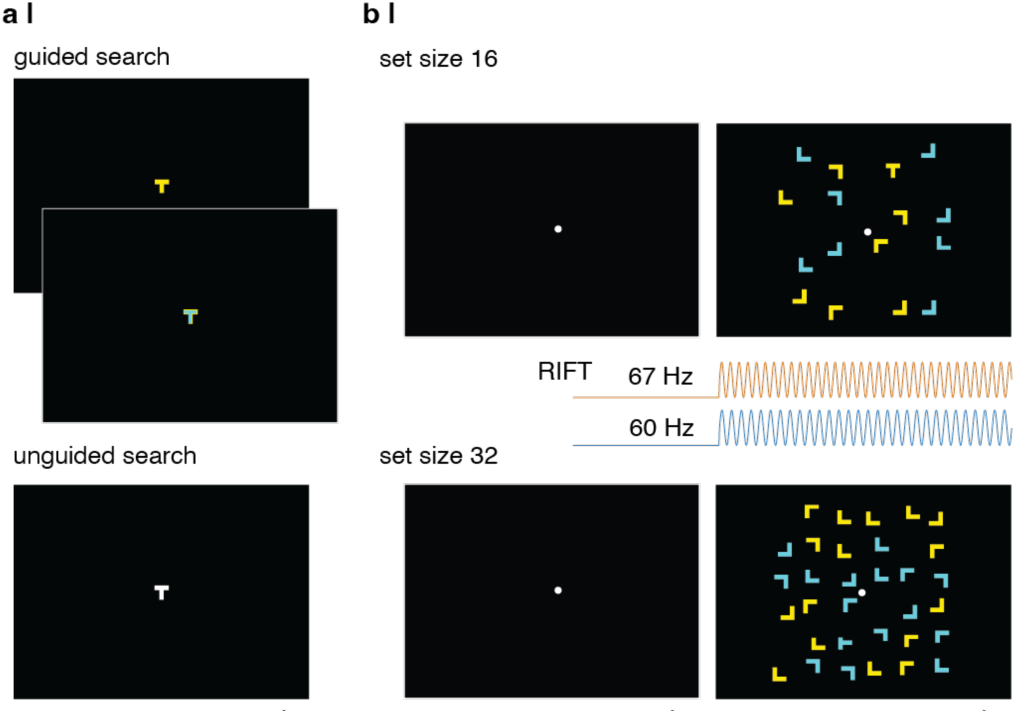
Experimental paradigm (compare Duecker et al., 2025). a| A colour cue at the beginning of a block indicated the target colour for the following 40 trials (guided search) or signalled the beginning of an unguided search block during which the colour of the target was randomized. b| Participants were asked to indicate whether a letter T was presented among 16 or 32 items. The number of items was kept constant during the block. The search stimuli were tagged, according to their colour, at 60 and 67 Hz. Tagging frequencies were randomized over trials.

### High alpha power prior to the search is associated with fast response times

We used a General Linear Model (GLM) spectrum approach (Quinn et al., 2024), with spectral magnitude as the dependent variable, to test whether power in the pre-search interval is correlated with reaction time (Supplementary Figure 1a). The Short time Fourier spectrogram for each participant, trial, and MEG sensor was estimated using a sliding window approach (Supplementary Figure 1a). The design matrix of the GLMs contained the intercept and the factors guided vs unguided, set size 32 vs 16, target present vs absent, time-on-task, distance to the parietal sensors (indicating slouching in the chair over the duration of the experiment), and reaction time. A GLM with these factors was then fitted to the magnitude at each channel (combined planar gradiometers), frequency, and timepoint, separately per participant (Supplementary Figure 1b, Griffiths et al., 2021; Quinn et al., 2024).

To account for spurious correlations between the auto-correlated reaction time and alpha power, we generated a null-distribution by randomizing over blocks (Harris, 2021, Supplementary Figure 1c, 500 permutations). The reaction time regressor was then z-scored using the mean and standard deviation of this empirically obtained null distribution before comparing all values to zero using a cluster-based dependent-sample t-test (over the -1 to 0s baseline interval, 5,000 permutations, see Methods and Supplementary Figure 1c). The topographies and time-frequency representations of all regressors except reaction time and time-on-task are depicted in Supplementary Figure 1d.

Figure 2a depicts the results of the cluster-based dependent sample t-test of the reaction time regressor, indicating a significant negative correlation between spectral power in the baseline interval and reaction time. The outline of the cluster suggests that the effect was driven by the alpha-band. Indeed, the projected spectra demonstrate that fast responses are associated with higher power at 10-14 Hz (Figure 2b). The spectra were predicted based on the GLM intercept and reaction time regressor applied to the maximum and minimum reaction time for each participant and normalized to the maximum power in the spectrum of the slow trial before averaging (also see Supplementary Figure 2 for the projected spectra for each participant). Importantly, comparison of the hit rates, i.e. a correct indication of target present vs absent, for trials with high and low alpha power, did not reveal any significant difference (dependent sample t-test t(30) =-0.16, p =0.87, CI=[-0.0096, 0.0082], Figure 2c, trials binned using a balanced median split approach as detailed below). This demonstrates that the speeded-up responses reflect an improvement in behaviour with higher alpha power.

In line with these findings, Supplementary Figure 1e, depicting the time-frequency representation of Cohen’s F^2^, demonstrates that including the reaction time regressor into the model explains larger portion of the variance for frequencies in the alpha-(8-12 Hz) and beta-band (15-25 Hz). A jackknife approach applied to the reaction time regressor revealed that the observed negative correlation between alpha power and reaction time was constrained to the -0.3s to 0s interval in the 10-14Hz band (Supplementary Figure 1f).

Figure 2d shows the topography and time-frequency representation of the time-on-task (tot) regressor, indicating an increase of alpha power with task duration (T-values obtained by normalizing the regressors by the variance of the residuals). This regressor is linearly separable from reaction time, as both factors were included in the same multiple regression model. Importantly, comparison of Figure 2a and d suggests differences in the spatial and spectral properties of the correlations between alpha and reaction time and alpha and time-on-task.

In sum, the GLM-spectrum approach supports our hypothesis that strong alpha oscillations in preparation for the search are associated with faster responses in visual search. These results were not attributable to practice effects or increased fatigue with task duration.

**Figure 2.**
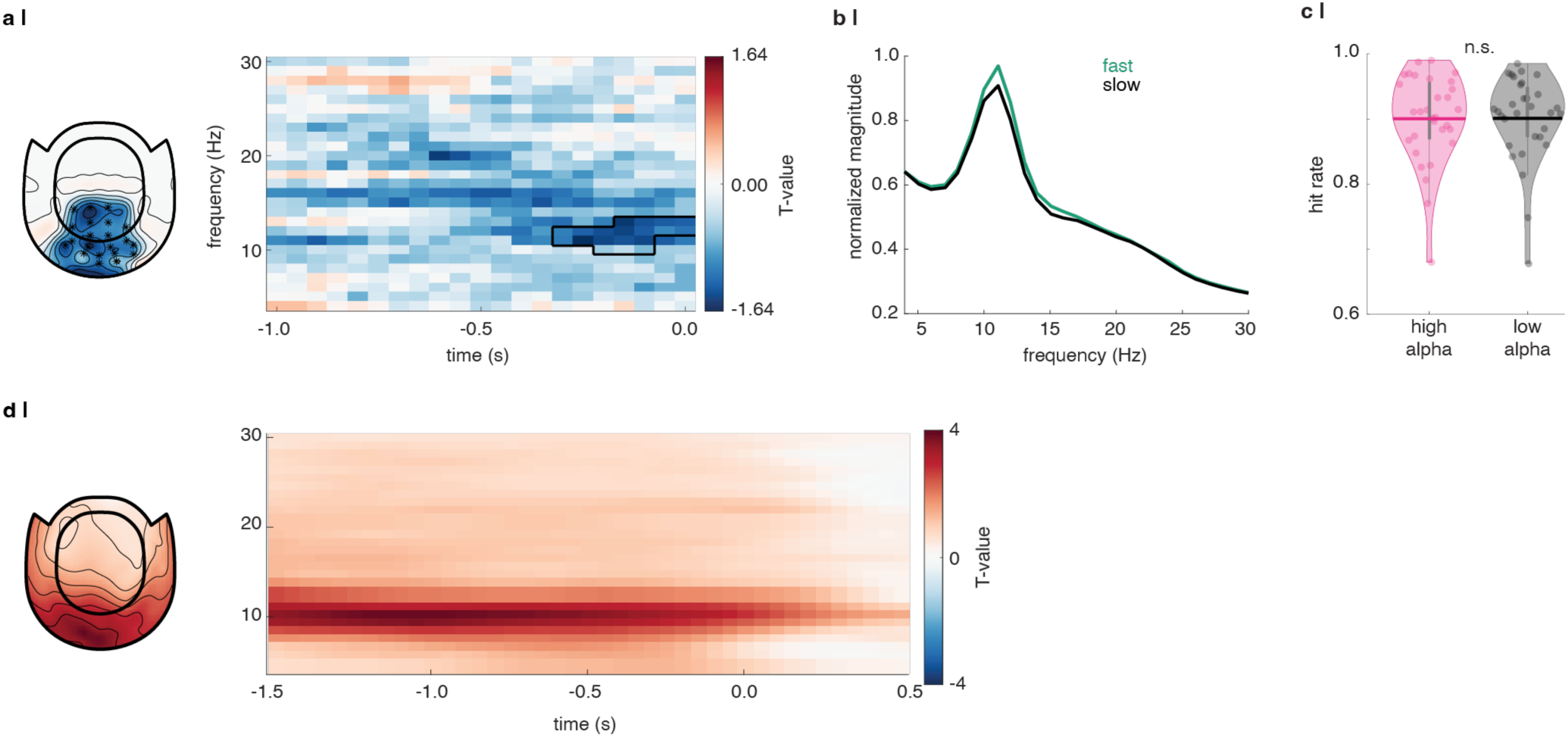
The GLM approach reveals a significant negative correlation between alpha power and reaction time when accounting for time-on-task **a |** Negative correlation between pre-search alpha power and reaction time. (left) topography of the T-statistics of the cluster-based permutation test comparing the reaction time regressor to zero, averaged over the -0.3 to 0 s interval, in the 10-14Hz band. Frontal and temporal sensors were not included in the cluster-based test and have therefore been set to 0. Asterisks indicate all sensors included in the cluster. (right) Time-frequency representation of the T-values, averaged over the sensors marked in the topography. The outline marks the extent of the cluster. **b |** Projected spectra were obtained for each participant by applying the reaction time regressor to the maximum and minimum reaction time and adding it to the estimated intercept. The spectra were normalized to the maximum power value in the “slow” spectrum for each participant before averaging. Fast reaction times are associated with higher power in the alpha-band. **c |** Hit rate (correct identification of target absent or present) does not significantly differ between high and low alpha trials (balanced median split accounting for time-on-task, t(30) =-0.16, p =0.87, CI=[-0.0096, 0.0082]). **d |** The time-on-task (tot) regressor indicating a strong positive correlation between alpha power and task duration. Note that the T-values were calculated by normalizing the regressors in the GLM, and are not the result of a cluster-based test.

### Relationship between alpha power and Rapid Invisible Frequency Tagging responses not established

We hypothesised that the correlation between high alpha power and reaction time underlies an inhibition of either the distractor stimuli or an inhibitory threshold applied to all stimuli. To this end, we correlated the RIFT signal with the average power at 10-14 Hz in the sensors with reaction time z-score < -1.96 for each participant. To account for confounds of time-on-task when correlating alpha power and RIFT responses, we used a balanced median split approach, applied to the magnitude-squared coherence, as well as a GLM fitted to the single-trial RIFT responses (Duecker et al., 2025). The balanced median split approach did not reveal any evidence for significantly reduced RIFT responses in the set size 16 (Figure 3a) or set size 32 (Figure 3b) search condition (no cluster formed, p > 0.5). Similarly, the GLM fitted to the single trial RIFT response in each sensor as a function of guided vs unguided, alpha power and time-on-task did not reveal any evidence that the RIFT response is reduced in trials with high pre-search alpha power. This was the case when fitting the GLM to the average RIFT response to stimuli in the target and distractor colour in guided and unguided search (Figure 3c, top), for the average RIFT response in the guided search condition (Figure 3c, middle), or for the response to the distractor in the guided search condition (Figure 3c, bottom).

In sum, while the negative correlation between alpha power and reaction time established above supports our hypothesis that alpha oscillations may facilitate the search, we were unable to find evidence for either of the hypothesised inhibitory mechanisms.

**Figure 3.**
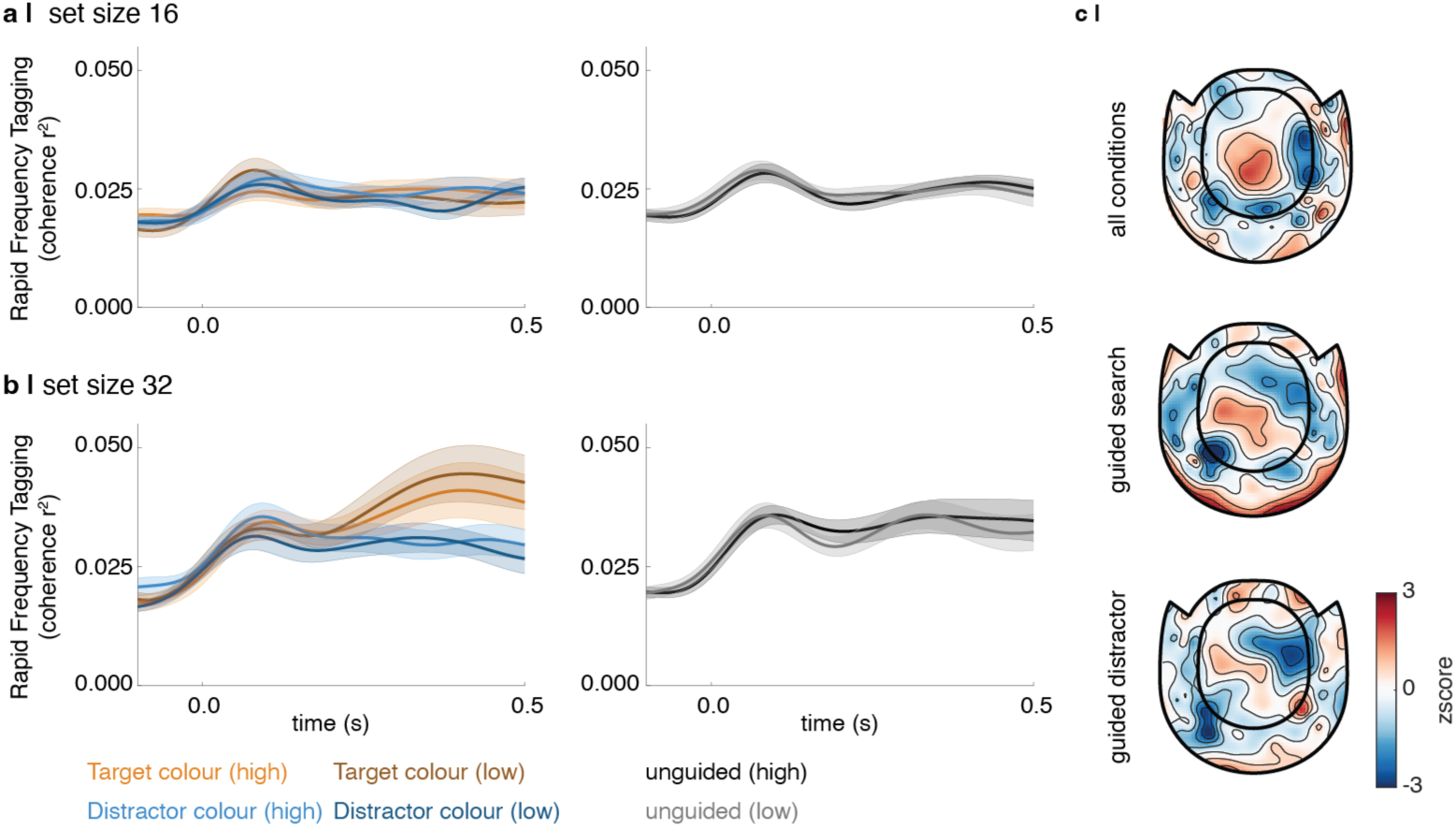
Correlation between alpha power and Rapid Invisible Frequency Tagging (RIFT) could not be established. **a,b |** Balanced median split on the magnitude-squared coherence reveals no difference in the RIFT responses for high and low alpha trials for either set size. **c |** GLM applied to the single trial RIFT responses. (top) A GLM fitted to the average RIFT response to stimuli in the target and distractor colour, with the factors set size, guided vs unguided, alpha power and time-on-task, did not reveal a significant correlation between alpha and RIFT (corrected for spurious correlations, 1,000 permutations, significance tested using cluster-based dependent sample t-test). (middle) Similarly, a GLM with the factors set size, alpha power and time-on-task fitted to the average RIFT response and a GLM fitted to the distractor stimuli in the guided search condition (bottom), did not reveal any evidence for a reduced RIFT response with high alpha power.

## Discussion

In this study, we combined MEG with RIFT in a classic visual search task to investigate whether alpha oscillations support visual search through inhibition. Our results demonstrate that high alpha power in the pre-search interval is correlated with faster response times. We hypothesised that the alpha oscillations modulate the gain of the visual inputs to either selectively inhibit distractors or apply an inhibitory threshold to all stimuli (Ph.D. thesis Pastuszak, 2018). However, we did not find evidence for a significant correlation between pre-search alpha power and the RIFT response. While our results suggest that enhanced alpha oscillations support visual search, evidence does not support that this is accomplished via gain modulation (Antonov et al., 2020; Gundlach et al., 2020; Zhigalov & Jensen, 2020).

Our results link alpha oscillations to enhanced performance in complex visual search tasks that require the suppression of a high number of distractors. This finding is surprising as it contradicts associations of alpha oscillations with cortical idling, drowsiness, and boredom (Adrian & Matthews, 1934; Pfurtscheller et al., 1996), as well as more recent work linking increased alpha power to reduced visual performance (Dijk et al., 2008; Hanslmayr et al., 2007; Iemi et al., 2017). As previous work linking alpha oscillations to mental fatigue has reported a drop in visual search performance (Fan et al., 2015; Hanna et al., 2018), we argue that our results show a functional role of alpha oscillations in visual search.

Several direct tests of whether alpha oscillations support distractor suppression in visual tasks have led to inconclusive results (Antonov et al., 2020; Forschack et al., 2022; Gundlach et al., 2020; Zhigalov & Jensen, 2020). While spatial attention tasks have repeatedly revealed an increase in alpha power over cortical regions processing distracting stimuli, other reports question to what extent the modulation of alpha power reflects an active mechanism that facilitates attention (reviewed in Morrow et al., 2023). A recent revised account based on novel evidence (Gutteling et al., 2022) argues that instead of directly suppressing distractors, alpha oscillations may be modulated by a mechanism controlling the allocation of attentional resources based on perceptual target load (Jensen, 2024). According to this view, alpha oscillations may operate in higher-order visual regions, and not the early visual cortex responding to the RIFT signal (Zhigalov & Jensen, 2020). While the link between alpha power and behaviour demonstrated in this and other studies indicates a functional role of alpha in visual tasks, it also supports the view that alpha oscillations may not suppress distracting information directly.

Considering these findings, we argue that alpha oscillations may not support the perceptual selection of items in visual search but instead aid the retention of the search template in visual working memory (VWM). Numerous studies have revealed how items retained in VWM affect visual search performance and the efficacy of distractors to capture the participant’s attention (reviewed in Huynh Cong & Kerzel, 2021; Olivers et al., 2011; Wolfe, 2021). Access to search templates in visual search has further been shown to require internal attention (Gazzaley & Nobre, 2012), to filter the items in VWM (Awh & Vogel, 2008) and to assign resources to each item based on task-relevance (Franconeri et al., 2013). Both internal attention (Foxe et al., 1998) and VWM (Bonnefond & Jensen, 2012; Fan et al., 2015; Jensen et al., 2002; Klimesch, 2012; Wianda & Ross, 2019; Zhou et al., 2023) have been associated with an increase in alpha power. Moreover, recent work using transcranial alternating current stimulation suggests that shifting the phase of ongoing alpha oscillations perturbs working memory performance (Chen et al., 2023). The idea that the relationship between alpha power and reaction reported here underlies VWM could be tested either with a multi-template search task (Grubert et al., 2025) or by using a complex, detailed image as the search target (Ackermann & Landy, 2014). We predict that alpha power will be modulated by the working memory load associated with the target(s).

Recent work has cautioned about the confounding relationship between alpha power and task duration (Benwell et al., 2019), demonstrating that failure to account for time-on-task may result in spurious correlations between alpha power and performance (Kopčanová et al., 2024). Our results demonstrate a robust relationship between alpha power and reaction time when controlling for this confound. Furthermore, the time-frequency representation and topographies of the reaction time and time-on-task regressor suggest differences in the peak frequency and cortical region of the two regressors. While the GLM approach does not allow to disentangle the different sources of the alpha rhythm, a rich body of literature supports the idea that alpha oscillations emerge from different cortical circuits. For instance, separable alpha rhythm generators have been identified based on intracranial work in non-human primates (Bollimunta et al., 2008, 2011) and MEG studies in human participants (Giehl & Siegel, 2024). These results are supported by electrophysiological findings and computational models, that have identified different generators of the alpha rhythm, including a thalamic pacemaker (Becker et al., 2015; Bollimunta et al., 2011; Hughes & Crunelli, 2005; Vijayan & Kopell, 2012) and cortical circuits (Lopes da Silva, 1991a, 1991b, 1991c; Lopes Da Silva & Storm Van Leeuwen, 1977) driven by layer 5 pyramidal neurons (Jones et al., 2000), as well as traveling waves propagating between sensory and frontal areas (Alamia et al., 2023; Bahramisharif et al., 2013; Davis et al., 2020; Zhang et al., 2018). Different generators of alpha oscillations have further been suggested to serve different purposes in attention and working memory tasks (Rassi et al., 2024; Rodriguez-Larios et al., 2022; Sokoliuk et al., 2019; Zhou et al., 2025). We propose that the different time-frequency representations and topographies for the correlations between alpha power and reaction time and alpha power and time-on-task suggest the existence of different generators of alpha: one reflecting a mechanism supporting the task, and the other one reflecting idling or fatigue increasing with time. We believe that our findings pave the way for future investigations of the different cortical generators of these oscillations, as well as an understanding of alpha oscillations as a cortical mechanism for inhibition serving various purposes in cortical and thalamo-cortical networks.

## Conclusion

Our results demonstrate that enhanced occipital alpha power in preparation for a visual search task is associated with faster response times. In contrast to what we hypothesised, we were unable to establish a robust correlation between alpha power and the modulation of the RIFT responses to the search stimuli. While our findings suggest that high alpha power marks better performance, it remains to be tested if and how the rhythm causally influences search. We argue that the solution to this puzzle lies in the consideration of different cortical generators of alpha, as well as different mechanisms involved in visual search that may not directly be linked to gain control.

## Methods

The recruitment of participants (31 healthy volunteers), experimental design, and display physics are described in depth in a related publication (Duecker et al., 2025). Chiefly, the participants’ task was to indicate whether a cyan or yellow letter “T” was presented among “L”s, while the magnetoencephalogram was recorded (MEGIN Triux, MEGIN Oy, Espoo, Finland, Figure 1). A cue at the beginning of each block indicated whether the search was guided or unguided for the following 40 trials. In guided search, participants were informed about the colour of the “T” for the duration of the block. In the unguided search condition, the target colour was randomised over trials. The experiment consisted of a total of 24 blocks (960 trials). Rapid Invisible Frequency Tagging (RIFT) at 60 and 67Hz was applied using a Propixx lite projector (VPixx Technologies Inc, Quebec, Canada) set to a refresh rate of 480 Hz, to tag the stimuli according to their colour. Each search was preceded by a 1.5s baseline interval, during which we expected an increase in alpha power. All stimuli were created using the Psychophysics Toolbox version 3 (Brainard, 1997) in MATLAB 2017a (The Mathworks, Natick, MA, USA).

### MEG analysis

The pre-processing of the MEG data included a correction of the faulty MEG sensors, Signal Space Separation (“Maxfilter”), and semi-automatic artefact rejection using Independent Compontent Analysis. Additionally, we discarded trials with short response times (<0.2s) and without a response. The photodiode channels recording the RIFT signal were replaced by a perfect sine wave. The magnitude-squared coherence between the RIFT signal and each MEG sensor was calculated based on the bandpass-filtered, Hilbert-transformed signals as outlined in Duecker et al., 2025. The single trial RIFT response was calculated based on the Fast Fourier Transform with 100ms time windows, moved over the 0.2 to 0.5s interval with a 75% overlap. The significant RIFT sensors, shown for the magnitude-squared coherence results, were identified using a permutation-based approach as described in Maris & Oostenveld, 2007 (Duecker et al., 2025; Pan et al., 2021).

### The General Linear Model spectrum

The correlation between spectral power and reaction time, and RIFT response and alpha power were investigated using a General Linear Model spectrum approach (Quinn et al., 2024). The single-trial power at each sensor, frequency, and timepoint was modelled as a linear function with the regressors constant (intercept), guided vs unguided (coded as 1 and -1, respectively), set size 32 vs 16, target present vs absent, time-on-task, slouch, and reaction time (Supplementary Figure 1b). A “slouch” regressor, accounting for the participants sliding down in the chair and thus the MEG helmet over the duration of the experiment, was estimated based on the distance between the participants head and the parietal sensors, which was acquired every 10 minutes throughout the experiment. The beta regressors were estimated using the pseudoinverse of the design matrix (Quinn et al., 2024). The associated T-values were obtained by dividing the respective beta-value by the standard error of the residuals to create a test statistic for the hypothesis that the parameter estimates are different to zero. To account for spurious correlations based on the autocorrelations of the spectral power and reaction time distributions (Harris, 2021), we generated a null distribution by shuffling the order of the blocks and normalized the T-value of the reaction time regressor based on the mean and standard deviation of the empirically obtained distribution. The temporal and spectral extent of the cluster was determined using a jackknife approach. For each iteration, the cluster-based dependent sample t-test was applied to all but one participant, and the lower and upper time and frequency bound of the cluster were stored (Miller et al., 1998; Sassenhagen & Draschkow, 2019). The size of the scatters in Supplementary Figure 1c indicates the number of occurrences of the upper and lower time and frequency limit of the cluster.

### Statistical analyses of the relationship between alpha power and RIFT signal

We tested whether high alpha power is correlated with reduced RIFT responses using a balanced median split approach applied to the magnitude-squared coherence, as well as a GLM fitted to the single trial RIFT data.

For the balanced median split, we first sorted the single trial MEG data into four bins based on task duration (from early to late). In each bin and search condition, we split the data based on spectral power in the 10 to 14 Hz band in the combined planar gradiometers for which the z-score of the reaction time regressor in the GLM described above was z < -1.96. For each condition, we then calculated the coherence between the MEG sensors and RIFT signal separately for all trials for which alpha power was above and below the median. To test whether high alpha power was associated with overall reduced RIFT responses, we averaged the coherence time series for targets and distractors in the high alpha trials and compared those to the averaged coherence over both stimuli in the low alpha trials using a cluster-based permutation t-test (5,000 permutations).

The GLM was fitted to the single trial RIFT response, averaged over the combined planar gradiometers (see above) over trials, with the regressors guided vs unguided, time-on-task and alpha power at 10-14 Hz averaged over all sensors with a reaction time z-score < -1.96. The estimated alpha power coefficients were then normalized to a null distribution, obtained as described above, and compared to zero using a cluster-based permutation t-test (5,000 permutations).

The code for the experimental paradigm and data analyses are shared at https://github.com/katduecker/vs_rft.

## Supplementary Figures

**Supplementary Figure 1.**
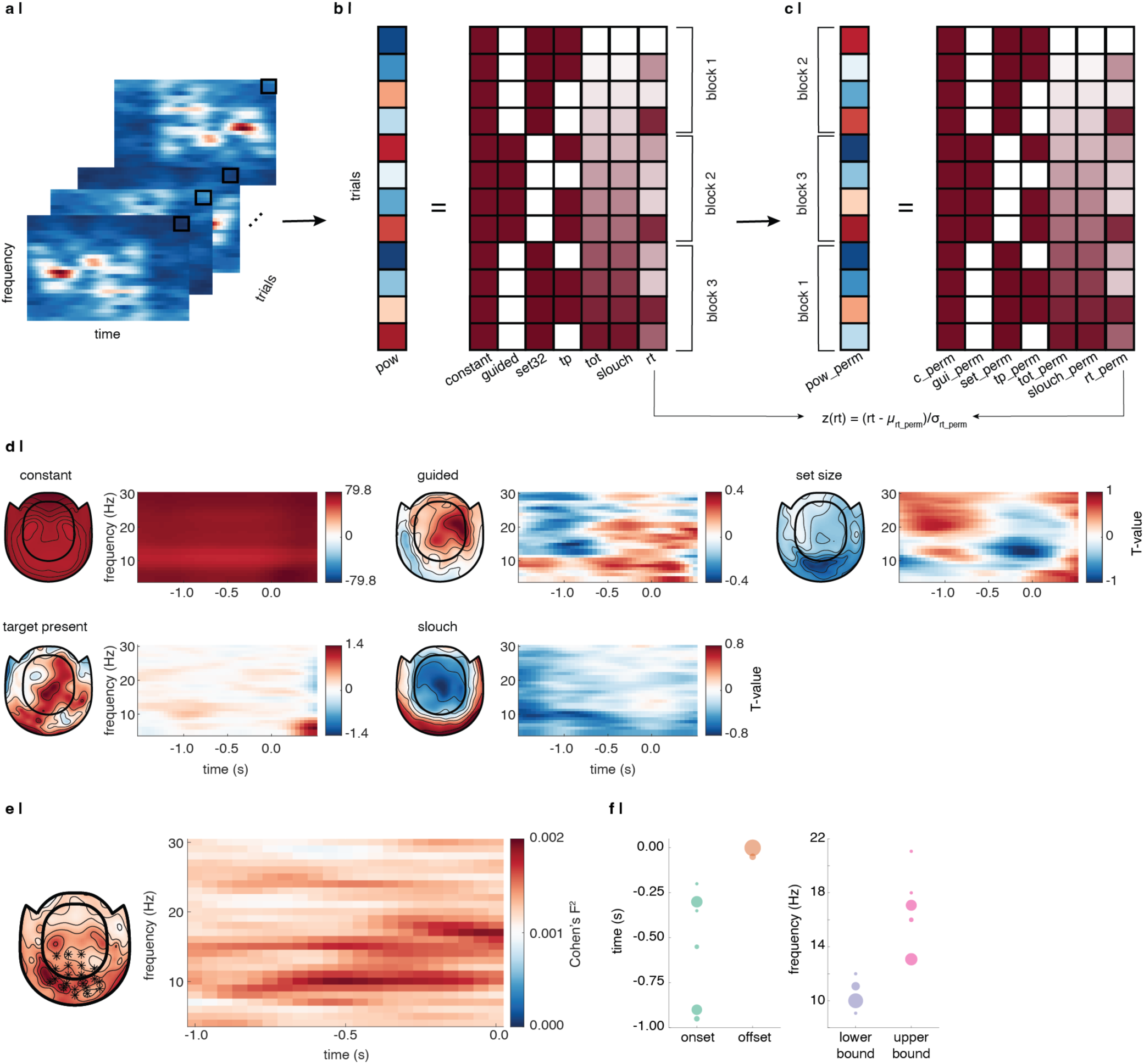
a | Single trial time-frequency representations of Fourier magnitude were estimated using a 1-second Hanning taper, moved over the time series in steps of 0.05s. **b |** The GLM spectrum was estimated by extracting the magnitude values at each sensor (combined planar gradiometers), frequency, and timepoint over all trials, and fitting a GLM with the regressors constant (intercept), guided vs unguided (coded as 1 and -1), set size 32 vs 16 (1 vs -1), target present vs absent (1 vs -1), time-on-task (ranging from 0 to 1), distance to the parietal sensors (slouch, 0 to 1), and reaction time. **c |** A null distribution for the reaction time regressor was estimated by shuffling over blocks. **d |** GLM regressors (all but reaction time and time-on-task which are shown in Figure 2). Topographies were averaged over the -0.3 to 0 s interval before the stimulus onset and the 10 to 14Hz band, time-frequency representations were averaged over the significant sensors identified for each participant. The topography of the slouch regressor indicates an increase in power in the lower sensors and a decrease in parietal sensors, reliably capturing the participants increasingly relaxed postures over the duration of the task. **e|** Cohen’s F^2^ reveals that by inclusion of the reaction time regressor, the model can explain more variance in the alpha and beta band. **f|** The temporal and spectral extent of the cluster were identified using a jackknife approach, suggesting that the reaction time effect was focused on power in the 10-14Hz band in the -0.3 to 0s interval.

**Supplementary Figure 2.**
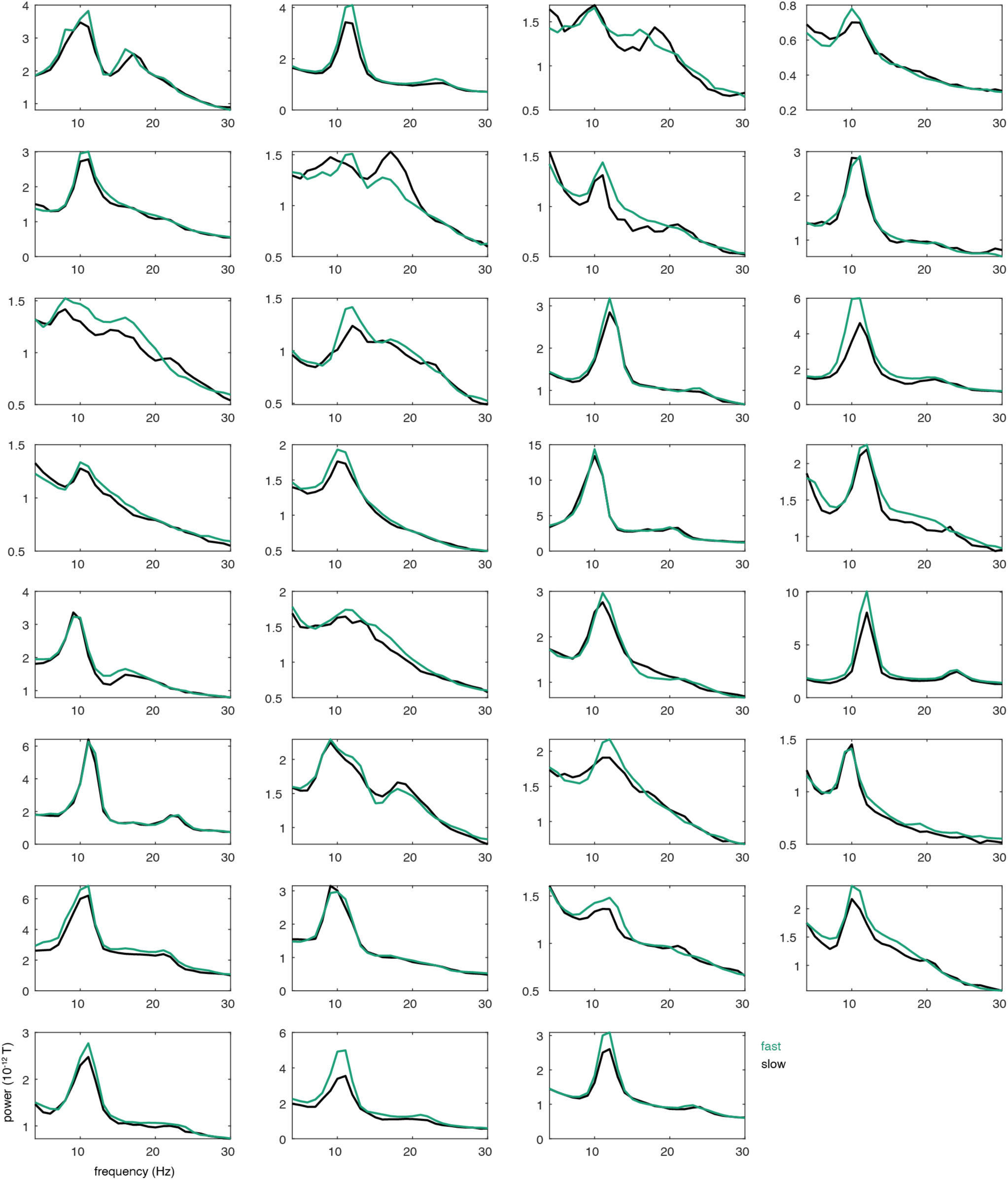
Projected spectra for fast vs. slow response times per participant.

## Author Contributions

A.P. conducted preliminary experiments and proposed blanket inhibition hypothesis, S.H. proposed research idea, K.D., K.L.S., S.H., J.W., and O.J. designed the experiment, K.D. acquired and analysed the data, Y.P., S.H. and O.J. supported analysis of MEG data, B.J.G. and A.J.Q. supported the development of the GLM analysis, K.D. and O.J. wrote the paper, all authors edited the paper.

## Acknowledgements

The authors thank Veikko Jousmaki for providing the light-to-voltage converter and Jonathan L. Winter for support with the MEG data collection.

## Grant support

This research was funded in whole, or in part, by the Wellcome Trust (227420) to O.J. Y.P. is supported by a Leverhulme Early Career Fellowship (ECF-2023-626). B.J.G. is funded by a Leverhulme Trust Early Career Fellowship (ECF-2021-628). J. W. is funded by NIH EY017001.

## Notes

### Competing Interest Statement

The authors have declared no competing interest.

